# Mathematical modelling of activation-induced heterogeneity reveals cell state transitions underpinning macrophage responses to LPS

**DOI:** 10.1101/2021.09.19.461010

**Authors:** S. Dey, D. Boucher, J. W. Pitchford, D. Lagos

## Abstract

Despite extensive work on macrophage heterogeneity, the mechanisms driving activation induced heterogeneity (AIH) in macrophages remain poorly understood. Here, we use two *in vitro* cellular models of LPS-induced tolerance (bone marrow-derived macrophages or BMDMs and RAW 264.7 cells), single-cell protein measurements, and mathematical modelling to explore how AIH underpins primary and secondary responses to LPS. We measure expression of TNF, IL-6, pro-IL-1β, and NOS2 and demonstrate that macrophage community AIH is dependent on LPS dose. We show that altered AIH kinetics in macrophages responding to a second LPS challenge underpin hypo-responsiveness to LPS. These empirical data can be explained by a mathematical 3-state model including negative, positive, and non-responsive states (NRS), but they are also compatible with a 4-state model that includes distinct reversibly NRS and non-responsive permanently states (NRPS). Our mathematical model, termed NoRM (Non-Responsive Macrophage) model identifies similarities and differences between BMDM and RAW 264.7 cell responses. In both cell types, transition rates between states in the NoRM model are distinct for each of the tested proteins and, crucially, macrophage hypo-responsiveness is underpinned by changes in transition rates to and from NRS. Overall, our findings provide support for a critical role for phenotypically negative macrophage populations as an active component of AIH and primary and secondary responses to LPS. This reveals unappreciated aspects of cellular ecology and community dynamics associated with LPS-driven training of macrophages.

## 1 Introduction

Variability in gene expression in eukaryotic cells is required to allow communities of cells to switch from homeostatic to inducible states while responding to external cues (Blake et al., 2006;Eldar and Elowitz, 2010). Genetically identical populations show considerable cell-to-cell variability, particularly of proteins that are stress induced (Bar-Even et al., 2006;Newman et al., 2006). Studies on heterogeneity have found that expression of housekeeping genes tends to be normally (or log normally) distributed in apparently homogeneous populations (Kumar et al., 2014;Klein et al., 2015) while a subset of genes displays increased cell-to-cell variability with a bi-modal distribution (Shalek et al., 2013). Population heterogeneity plays a critical role in shaping immune responses. For example, seemingly clonal populations of myeloid cells can produce effector cytokines heterogeneously. Several models of myeloid heterogeneity have been described, including bi-phasic transcription factor activation such as that of NF-kB and autocrine/paracrine effects of TNF or IL-1β in response to TLR stimulation (Burns et al., 1998;Han et al., 2002;Caldwell et al., 2014;Hayden and Ghosh, 2014) and recently shown to be partly dependent on intercellular desynchronization of molecular clock (Allen et al., 2019). Interestingly, macrophage hypo-responsiveness to secondary stimulation has been associated with a switch in phenotype wherein, by a combination of TLR4 attenuation, microRNA (miRNA)-mediated silencing expression, and chromatin modifications, macrophages lose their ability to make inflammatory proteins (Biswas and Lopez-Collazo, 2009;Netea et al., 2015;Seeley and Ghosh, 2017) with alterations in chromatin accessibility being a more permanent cause for this phenomenon. In addition, macrophage hypo-responsiveness is driven by effects only on some genes while expression of others remains unaffected (Foster et al., 2007). Despite the above insight, the effect of primary or repeated stimulation on macrophage population heterogeneity, termed here as activation induced heterogeneity (AIH) and the underpinning molecular mechanisms remain elusive.

Here, we describe and quantify AIH within macrophage communities using two simple cellular systems of primary and secondary LPS challenge and measuring expression of four pro-inflammatory proteins. We built a mathematical model to propose and explore theoretical cellular states underpinning the empirically observed consistency of macrophage communities. Our analyses reveal that transitions to and from phenotyplically negative or non-responding macrophage populations are critical detemrinants of macrophage AIH and responses to primary and secondary LPS challenge.

## Materials and Methods

### Animals

Female C57BL/6 CD45.2 mice were obtained from Charles River (UK). Animal care were regulated under the Animals (Scientific Procedures) Act 1986 (revised under European Directive 2010/63/EU). Mouse breeding was performed under a UK Home Office License (project licence number PPL 60/4377) with approval from the University of York Animal Welfare and Ethical Review Body.

### Cell culture

BMDMs were isolated from female C57BL/6 mice and differentiated in the presence of MCSF1 (50ng/ml) for 6 days and then frozen at −70°C. Frozen BMDM from half a mouse (1 tibia and 1 femur) were plated and cultured in 10 mL of macrophage media in 100cm petri dishes for 2-3 days in the presence of MCSF1 before plating them on 24 well plates for experiments.

Both RAW264.7 cells and BMDMs were cultured in Dulbecco’s Modified Eagle Medium (DMEM) supplemented with 1% streptomycin-penicillin mixture, 1% L-glutamine and 10% fetal calf serum (Hyclone). For experiments using BMDM, MCSF1 was added in the cell culture media and kept for the duration of the experiment.

RAW 264.7 cells, a monocyte/macrophage-like cells, originating from Abelson leukemia virus transformed cell line derived from BALB/c mice, were detached for passaging using 1x Trypsin-EDTA (Invitrogen) by incubating at 37°C for 10 minutes. Cells were detached completely by gently scraping with cell scraper with a cross-ribbed handle (VWR). Upon reaching 70-80% confluency, cells were harvested and plated in 24 well plates. BMDMs were detached by gentle pipetting up and down using ice cold 1X PBS (Gibco). Cells were centrifuged (1500RPM for RAW264.7 and 1300RPM for BMDMs) at room temperature for 5 minutes for the purposes of washing or re-suspending.

### LPS challenge

LPS from *Escherichia coli* serotype 055:B5 (Sigma-Aldrich, L2880) was used. This is a phenol extracted LPS with <3% protein impurity. 200-250,000 RAW264.7 cells or BMDMs were plated overnight before experiments. All cells were plated in a Corning 24 well plate in 500ul of DMEM. For LPS titration experiments, cells were either stimulated with LPS or were left in media (untreated) on day 1. Cells were challenged with 1, 10, 100 or 1000 ng/ml of LPS. Supernatant was collected at 24 hours and stored at −20°C. Cells were harvested for flow cytometry at 16 or 24 hours from LPS stimulus.

For inducing hypo-responsiveness, cells were either stimulated with 10 or 1000 ng/ml of LPS or left untreated in media on day 0. After 24 hours (day 1), cells were washed twice with PBS and replaced with media (Media/Media) or with media containing 1000 ng/ml of LPS (10/1000; 1000/1000 or Media/1000).

### Flow Cytometry

RAW264.7 cells of BMDMs were collected after washing in ice-cold PBS and then detaching the cells with Accutase (BioLegend). Prior to collection, cells were incubated in 10ug/ml of BFA (Brefeldin A, Sigma). BFA was added to the culture four hours prior to harvest for staining.

Cells and all reagents were maintained at 4°C throughout the intra-cellular staining protocol. Harvested cells were washed twice in PBS and re-suspended in approximately 50ul of PBS. Cells were stained with 100ul of 1:1000 Zombie Aqua live/dead stain (BioLegend) in PBS on ice for 8-10 minutes in the dark. F_c_ receptors were blocked with 5ul of 2mg/ml rat IgG for five minutes. Cells were fixed with BD Cytofix and permeabilized with BD Cytoperm. Intracellular staining was performed with the cocktail of antibodies made in permeability buffer. BV421-TNF (MP6-XT22; BioLegend), APC-IL6 (MP5-20F3; BioLegend, eFluor 610-NOS2 (CXNFT; ThermoFisher Scientific), PE-pro-IL-1β (NJTEN3, ThermoFisher Scientific), FITC-F4/80 (BM8, BioLegend) and PE-Cy7 CD11b (M1/70, BioLegend) were used for staining RAW264.7 cells. BMDMs and RAW264.7 cells were pre-gated on live cells, singlets, forward scatter, and side scatter (for gating intact cells), F4/80+ and/or CD11b for an average of 100,000 cells were collected per treatment.

### ELISA and Greiss Assay

IL-6, TNF and IL-1β concentrations in the cell culture supernatant were measured by enzyme-linked immunosorbent assay (ELISA) using BioLegend’s ELISA MAX Standard. Manufacturer’s recommended protocol was followed. Absorbance was read at 450nm with a wavelength correction at 570nm using a VersaMax Microplate Reader (Molecular Devices). Standard curves were generated using 4-parameter non-linear fitting to known standard concentrations using SoftMax Pro software. Optical density of the unknowns that fit within the linear range of the standard curve was used to calculate the concentration of the sample.

Greiss assay was used to measure nitrite concentrations in the supernatants. Diazotization reaction in Greiss assay was carried out as per manufacturer’s instructions (Promega). Plates were read on VersaMax microplate reader capturing absorbance between 520 and 550nm.

### Mathematical Modelling

Bespoke MATLAB code, NoRM, was written to implement stochastic simulations using the Doob-Gillespie algorithm. Parameter estimation of stochastic models were carried out by running the NoRM model with 10^5^-10^6^ sets of randomly generated parameter sets from a mixture of negative binomial, uniform, and normal distributions. The key transition rates (α, β, β_2_ γ_1_, γ_2_) were estimated using rejection sampling. The unitless μ (co-efficient for modelling LPS dynamics), was adjusted between the range of 0.1-100 to account for sensitivity to LPS for the four different proteins. The LPS decay rate, δ, was arbitrarily chosen and fixed at 0.5; model outcomes were qualitatively unaffected by this choice. Selection of parameters that explained empirical datasets was performed by rejection sampling based Akaike Information Criterion (AIC), with particular attention being paid to the key transition rates (α, β, β_2_ γ_1_, γ_2_). The modelling process is described in detail in the **Supplementary Material under “Supplementary text: Modelling Process”**.

### Statistics

All experiments were performed in at least three biological replicates. BMDM experiments were performed with macrophages from at least three mice. Statistical analysis was done using Graphpad Prism 6, Matlab and R. Treatment groups were compared using unpaired Student’s t-test.

## Results

### Macrophage community AIH is dependent on LPS dose

To capture distinct macrophage subpopulations upon activation with LPS we measured protein expression of three cytokines, TNF, pro-IL-1β and IL-6, and one intracellular pro-inflammatory protein NOS2, an enzyme that catalyzes nitric oxide formation. We selected these factors as they are all inducible upon LPS challenge. Furthermore, heterogeneity in TNF and IL-1β secretion in populations can be a result of bi-phasic NF-kB activation (Tay et al., 2010). IL-6 and NOS2 are also up-regulated due to LPS (Farlik et al., 2010;Tanabe et al., 2010). Also, all these proteins have been implicated in LPS-induced macrophage hypo-responsiveness. To study AIH, we focused on the early stages (within the first 24h) post-primary or secondary stimulation with LPS to minimise confounding effects of secondary and tertiary cytokine-mediated effects.

First, we selected RAW264.7 cells as a cellular model. These cells are thought to be a model of primary bone-marrow derived macrophages with regards to expression of surface receptors and the response to microbial ligands (Berghaus et al., 2010). We reasoned that using a macrophage cell line to study AIH also reduced the level of starting population heterogeneity in comparison to that we would observe using primary macrophages. The community composition of LPS stimulated RAW264.7 cells was represented graphically by charting the 16 possible sub-populations by adapting the Simplified Presentation of Incredibly Complex Evaluations (SPICE) method (Roederer et al., 2011), with each slice representing a subpopulation (**Figure 1A, B**) with positive fractions selected based on appropriate isotype controls (supplementary **Figure S1A**). Consistent with the concept of AIH, we found that the dose of LPS can have qualitative effects on the diversity of the response; quadruple positive and TNF negative triple positive (TNF-proIL1β+IL6+NOS2+) cells appear prominently at higher doses of LPS (100, 1000 ng/ml; **Figure 1B**), while quadruple negative (TNF-pro-IL1β-IL6-NOS2-) sub-populations and single positive cells for TNF (TNF+pro-IL1β-IL6-NOS2-) appear at lower doses (1, 10 ng/ml; **Figure 1B**). Despite heterogeneous compositions of low and high dose of LPS, double-positive TNF-proIL1β+IL6-NOS2+ cells were a part of all LPS doses with little variability (1, 10, 100, 1000 ng/ml; **Figure 1B**) suggesting the presence of sub-populations with differential dependence on the magnitude of LPS dose.

**Figure 1:**
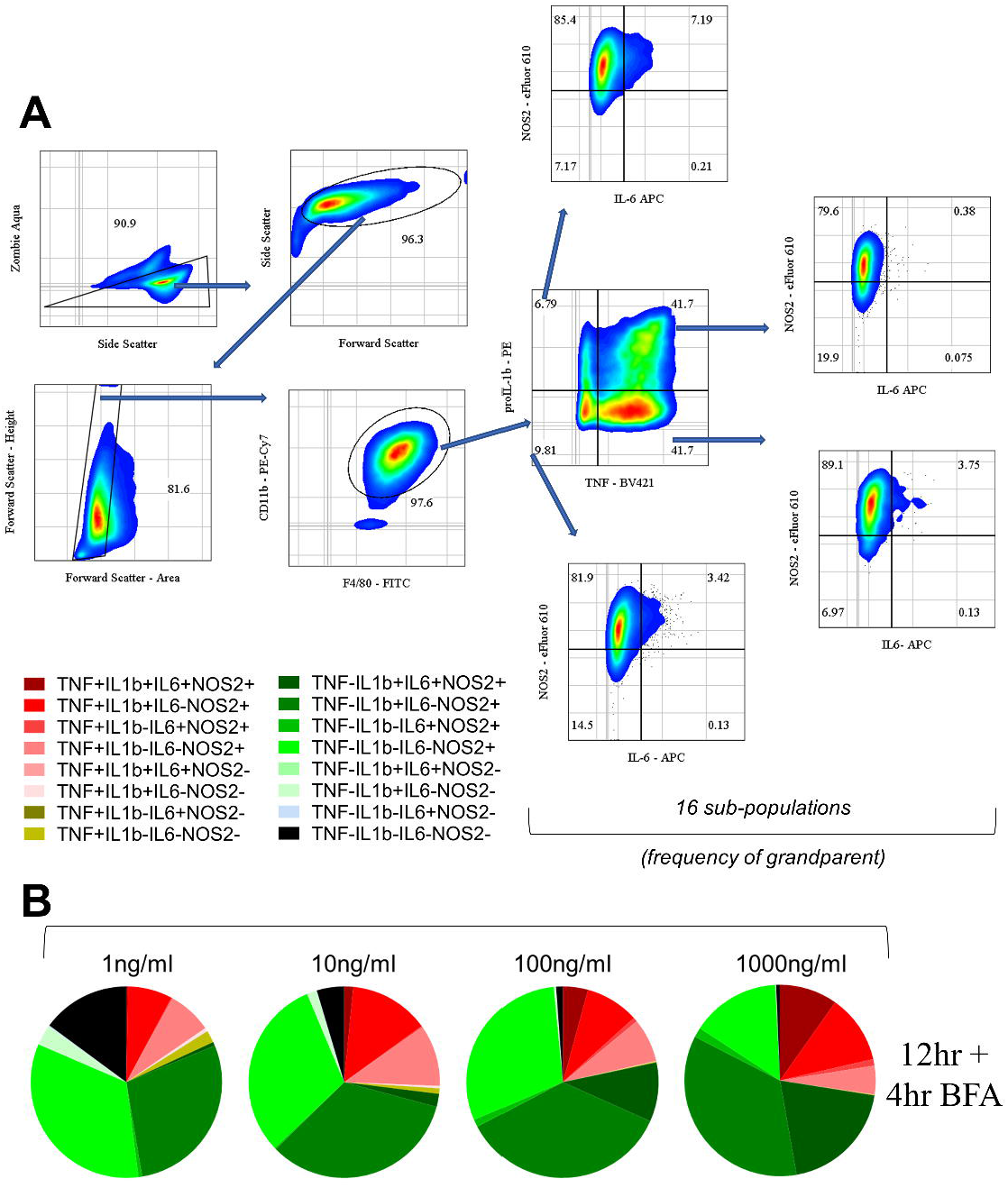
Macrophage community AIH is dependent on LPS dose. A. Flow cytometry gating to show 16 sub-populations determined based on TNF, IL-6, pro-IL-1b and NOS2. B. Pie charts represent the community composition at 16 hours post stimulus with the indicated doses of LPS for RAW264.7. Data representative of three independent experiments.

Next, we explored AIH in BMDMs, a cell model more faithfully capturing heterogeneity of primary macrophages. As in the case of RAW264.7 cells, exposure to LPS induced population heterogeneity in BMDMs albeit with different kinetics to that observed in RAW264.7 cells (compare **Figure 2** with **Figure 1B**). Whereas all populations were observed at 16h (12h stimulation followed by 4h of BFA treatment) post-stimulation in RAW264.7 cells, in BMDMs this was the case at earlier timepoints but not at 16h. Notably, the percentage of TNF-positive BMDMs peaked at 4h post stimulation, demonstrating a faster TNF response in BMDMs in comparison to RAW264.7 cells. In BMDMs, single positive cells for NOS2+ cells (TNF-pro-IL1β-IL6-NOS2+) increased while single positive cells for pro-IL1β (TNF-pro-IL1β+IL6-NOS2-) decreased with increasing magnitude of LPS dose at all time points (**Figure 2**). Further, quadruple negative sub-population (TNF-pro-IL1β-IL6-NOS2-) did not show a clear increase with a lower LPS dose as in RAW264.7 cells suggesting that the appearance of these sub-populations is more nuanced in BMDMs. While higher frequency of quadruple negative cells in 1ng/ml versus 10ng/ml could reflect differences in responses to increasing amounts of LPS the increased quadruple negative sub-population frequency in 100 and 1000ng/ml concentration may be due to a fast response accompanied by an immediate switch to a non-responding phenotype.

**Figure 2:**
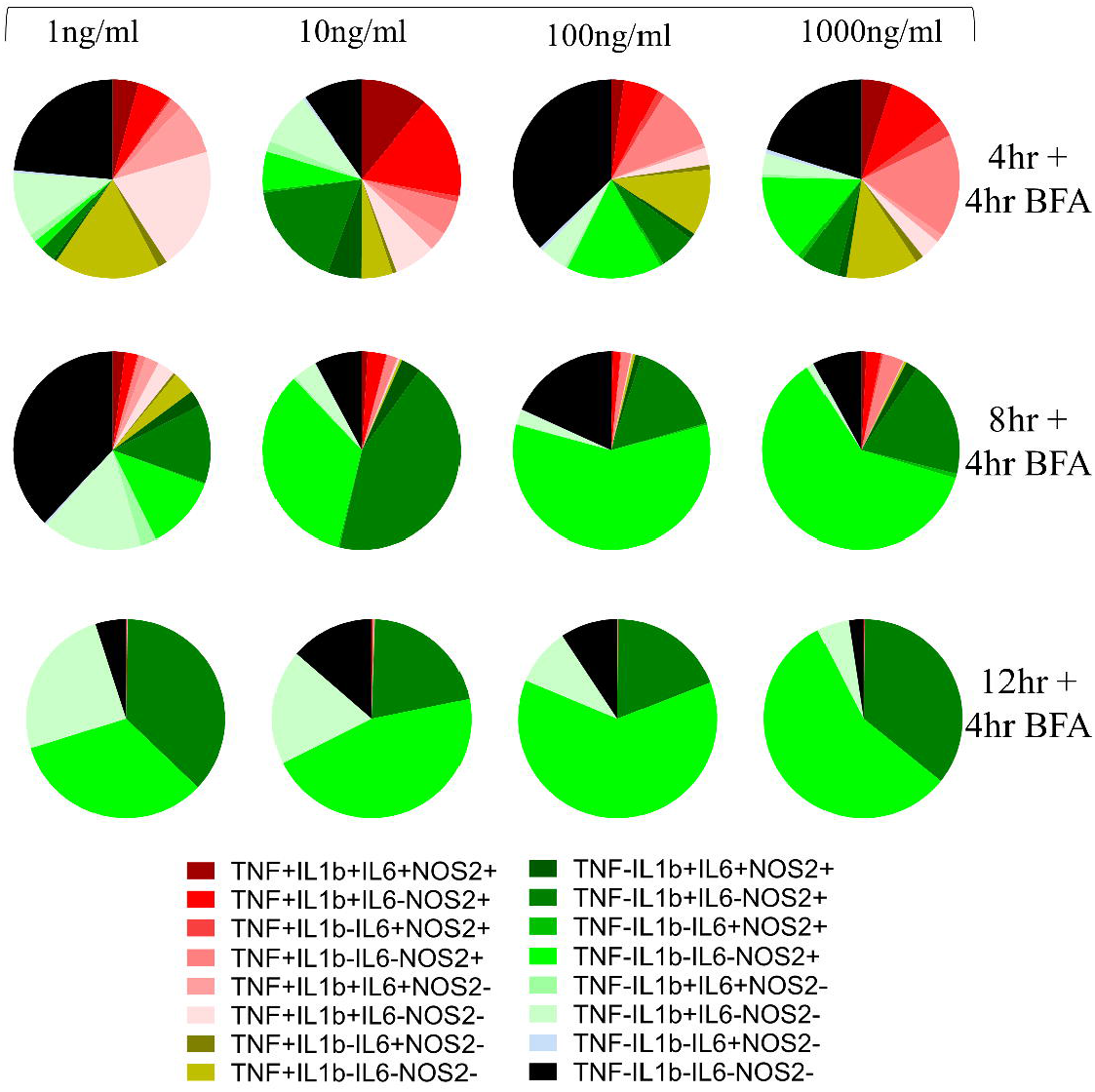
Macrophage community AIH kinetics for BMDMs. Pie charts represent the community composition at 8, 12 and 16 hours post stimulus with the indicated doses of LPS for BMDMs. BMDMs are pre-gated on Live/Singlets/FSC-SSC/CD11b+F4/80+ population. Data representative of three independent experiments.

Overall, our findings indicated that exposure to LPS induced population heterogeneity in macrophage communities for both a macrophage cell line (RAW264.7 cells) and primary BMDMs. As expected, cell-type specific differences were observed with BMDM responses occurring and peaking faster and reaching a plateau at lower LPS concentrations. These could be linked to differential sensitivity to LPS, but also differential pre-existing population heterogeneity between BMDMs and RAW264.7 cells. Regardless of these differences in kinetics our findings demonstrated that upon primary LPS challenge, AIH occurs in macrophages in an LPS-dependent manner.

### Altered AIH kinetics in response to a second LPS challenge underpin macrophage hypo-responsiveness

Next, we tested how changes in macrophage community compositions in RAW264.7 cells compare between macrophages challenged with LPS for a second time and macrophages responding to a first LPS stimulus. We obtained temporal snapshots of RAW264.7 cell communities responding to LPS alongside LPS responses of communities that were pre-exposed to varying LPS doses (**Figure 3A and Supplementary Figure 1B**). At the population level, cumulative secreted levels of TNF, IL-6, and NO were reduced for RAW264.7 cells (**Supplementary Figure 1C**), supporting that the LPS pre-treatment compromised the ability of cells to respond to a second LPS challenge. At the community level, single-cell measurements revealed that pre-treated macrophage community consistency (10/1000 and 1000/1000) differed to that seen during primary challenge (Media/1000) at 8h and 12h post stimulation but not at 16h (**Figure 3A**). For example, pre-treated macrophages were characterised by a prominent TNF-pro-IL1β+IL6-NOS2+ population but reduced TNF+ populations at the earlier stages of the response. This suggested that LPS-induced hypo-responsiveness is underpinned by different starting community compositions and altered community evolution trajectories, but not a different endpoint community composition.

**Figure 3:**
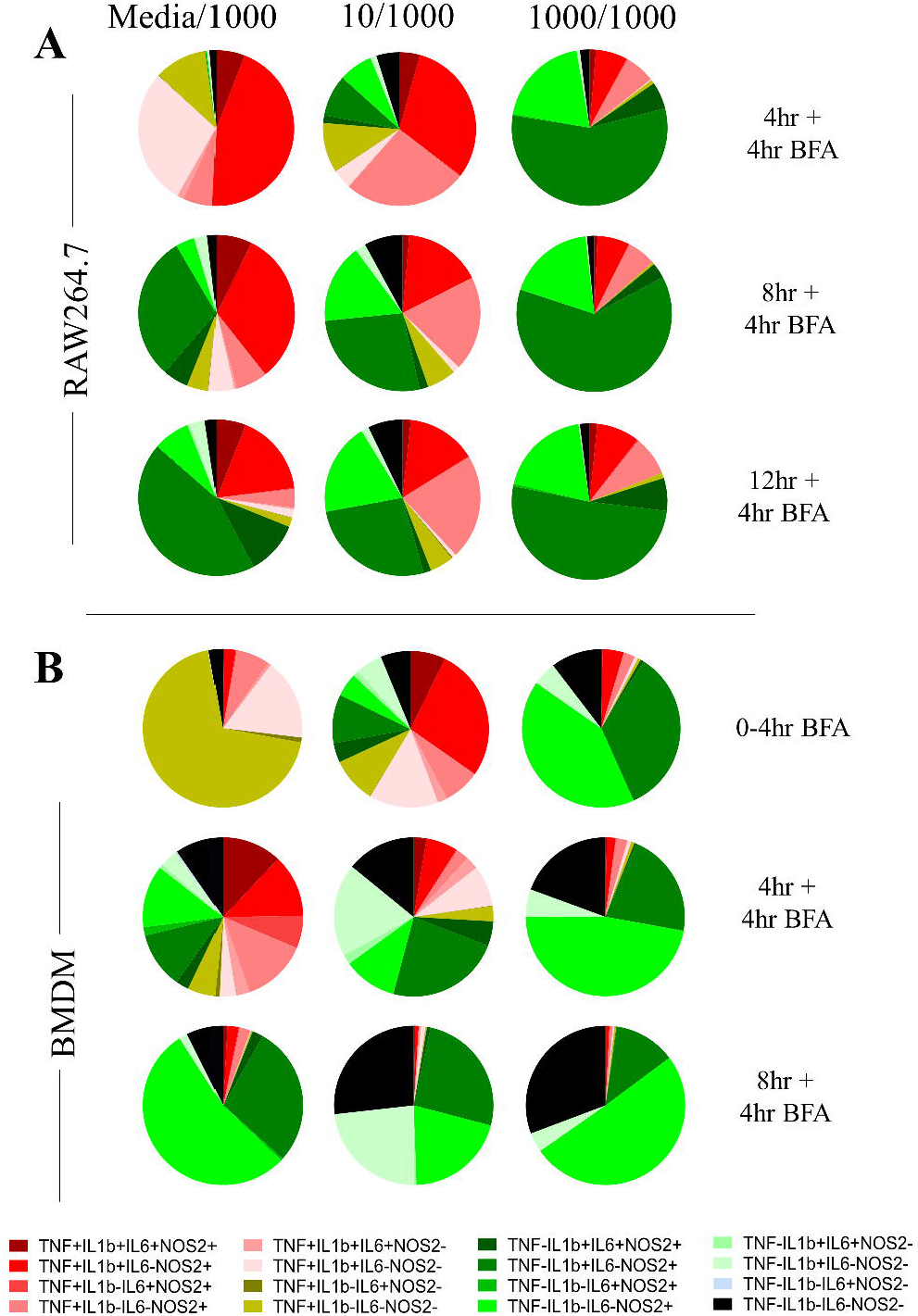
Altered AIH kinetics in macrophages responding to a second LPS challenge correlate with hypo-responsiveness. A. Averaged pie charts representing 3 independent experiments at the indicated timepoints post LPS challenge (1000ng/ml) of RAW264.7 macrophages pre-treated for 24 hours with either media (Media/1000), or 10ng/ml LPS (10/1000), or 1000ng/ml LPS (1000/1000). Legend indicates expression status for TNF, IL-6, pro-IL-1b and NOS2 subset. B. As in A, but for BMDMs.

In BMDMs, LPS-induced hypo-responsiveness was observed for cells pre-treated with 1000ng LPS for cumulative secreted levels of TNF, IL6, NO, and IL1β (**Figure S2A**). IL1β secretion was only observed in BMDMs pre-treated with 10ng LPS, in agreement with the known requirement of a priming step for pro-IL1β processing and IL1β secretion (Eder, 2009;Lopez-Castejon and Brough, 2011). BMDMs pre-treated with 1000ng LPS failed to produce secreted IL1β (**Figure S2A**), further supporting their hypo-responsive phenotype. At the single-cell level, despite increased levels at early timepoints (0-4hr) for NOS2 and pro-IL1β, we observed reduced expression of all measured proteins at 12 hours post stimulation of BMDMs pre-treated with 1000ng LPS (**Supplementary Figure S2B, C**) and increased quadruple negative population at all timepoints (**Figure 3B**). Having observed the kinetic differences between RAW264.7 cells and BMDMs upon primary LPS challenge (**Figures 1** and **2**), we explored an earlier time point in BMDM response (0-4hr in BFA). Indeed, 93% of all BMDMs also undergo an TNF+ state after which a fraction continues to be in the TNF+ sub-populations and a fraction that switches off (0-4hr BFA versus 4hr +4hr BFA in Media/1000 group). This finding is also in line with TNF being an early response protein (Bradley, 2008) and shaping macrophage community structure (Caldwell et al., 2014). As in the case of RAW264.7 cells, we observed more striking community differences during the early timepoints of the response (4hr + 4hr BFA and 8h + 4hr BFA) between pre-treated and control BMDMs. The end-point compositions (8h + 4hr BFA) were less distinct, although we note that in BMDMs, LPS pre-treatment resulted in an increase in quadruple negative cell populations and a reduction in TNF+ populations in LPS-pretreated cells at 12h post challenge (**Figure 3B**).

Interestingly, in the 10/1000 community 76% RAW264.7 cells (4hr+4hr BFA, **Figure 3A**; 68% BMDMs at 0-4hr, **Figure 3B**) were positive for TNF and hypo-responsiveness was most pronounced in the 1000/1000 community with just 8% BMDMs and 18% RAW264.7 cells being TNF+ in the first four and eight hours of the response respectively (**Figure 3**). Furthermore, in both BMDMs and RAW264.7 cells and at the earliest timepoints, the 10/1000 community showed a higher percentage of TNF+ cells than the 1000/1000 community that switch off rapidly to 45% for RAW264.7 cells and 26% for BMDMs at 12 and 8 hours respectively suggesting that a lower dose pre-stimulus decreases the capability of a population of cells to switch on TNF in response to a higher dose pre-stimulus. Furthermore, RAW264.7 communities of 10/1000 and 1000/1000 comprised of 5% and 2% negative sub-population respectively, confirming again that a small percentage of cells do not respond to the second dose of LPS. In addition, while overall TNF+ cells decrease over 8, 12 and 16 hours post LPS stimulus, the numbers of overall TNF+ cells first decrease (between 8 and 12 hours) then increase (between 12 and 16 hours) in RAW264.7 1000/1000 communities (**Figure 3A**). This suggested that a subset of cells can become positive for TNF later in response to the secondary stimulus.

Interestingly, in BMDMs it is the single positive NOS2 sub-population (TNF-pro-IL1β-IL6-NOS2+) and the double positive sub-population TNF-pro-IL1β+IL6-NOS2+ that dominates (41% and 34% respectively, **Figure 3B 0-**4hr) the first 4 hours of response in the 1000/1000 community compared with the Media/1000 community (**Figure 3B** 4hr). Similar to BMDMs, in RAW264.7 cells, the 1000/1000 community response in the first 8 hours also comprised of single positive NOS2 (20%, 4+4hr BFA **Figure 3A**) and double positive TNF-pro-IL1β+IL6-NOS2+ (57%, 4+4hr BFA **Figure 3A**). IL-6+ sub-populations were consistently reduced at all timepoints when compared between the Media/1000 and the 1000/1000 communities (**Figure 3B, Supplementary Figure S2**).

Overall, these results demonstrated that for both RAW264.7 cells and BMDMs, pre-exposure of macrophages to low or high LPS doses resulted in altered AIH kinetics during a secondary LPS challenge in comparison to macrophages receiving a primary LPS challenge. Endpoint community compositions showed modest differences between cells responding to one or two LPS challenges. The observed secreted protein hypo-responsiveness phenotype was predominantly reflected in the altered kinetics of changes in community composition upon LPS stimulation. Our data indicated that a critical part of the community response to LPS occured in the first 8-12 hours for RAW264.7 cells and 4-8 hours for BMDMs of the primary challenge, suggesting that at later time points a proportion of cells might be non-responsive in a reversible or permanent manner. We note that the effects were different for each of the measured proteins (**Figure 3**), suggesting that protein-specific mechanisms were involved in LPS-induced hypo-responsiveness.

### Transitions between distinct non-responding macrophage subsets underpin responses to LPS

To complement our empirical studies and understand how AIH contributes towards macrophage responses to LPS, we constructed conceptual mathematical models. At the heart of these models is the idea that any individual cell may make a transition from a non-protein producing state to a protein producing state (termed “negative” and “positive” states hereafter). These transitions occur at random in continuous time, and the probability (per unit time) of transition depends on the current environment of a cell such as the presence or absence of antigen (**Figure 4A, Supplementary text: Modelling Process**). The simplest models restricted each cell to be in a negative or a positive state only. While these models were found to be useful to understand antigen (LPS) dependent switching on and off for each individual protein independently (Eq 7, **Supplementary text: Modelling Process**), they failed to describe the ability of a subset of cells to become hypo-responsive that was suggested by our empirical studies without explicitly changing the rate at which a negative population switched to positive (**Supplementary text: Modelling Process**). Therefore, we refined the model by allowing two further cell states which reflect the empirical observations. Explicitly, we allowed the possibility that a positive cell could switch to a third non-responsive state (NRS, **Figure 4B**), generating a 3-state model. In addition, we also explored the possibility that cells in the NRS may make one of two transitions, either to a fourth, non-responsive permanently state (NRPS, **Figure 4C**) or back to the negative cell state, generating a 4-state model. Models were implemented using the Doob-Gillespie algorithm and were checked for faithfulness to the mean-field solution (**Supplementary Figure S3A**).

**Figure 4:**
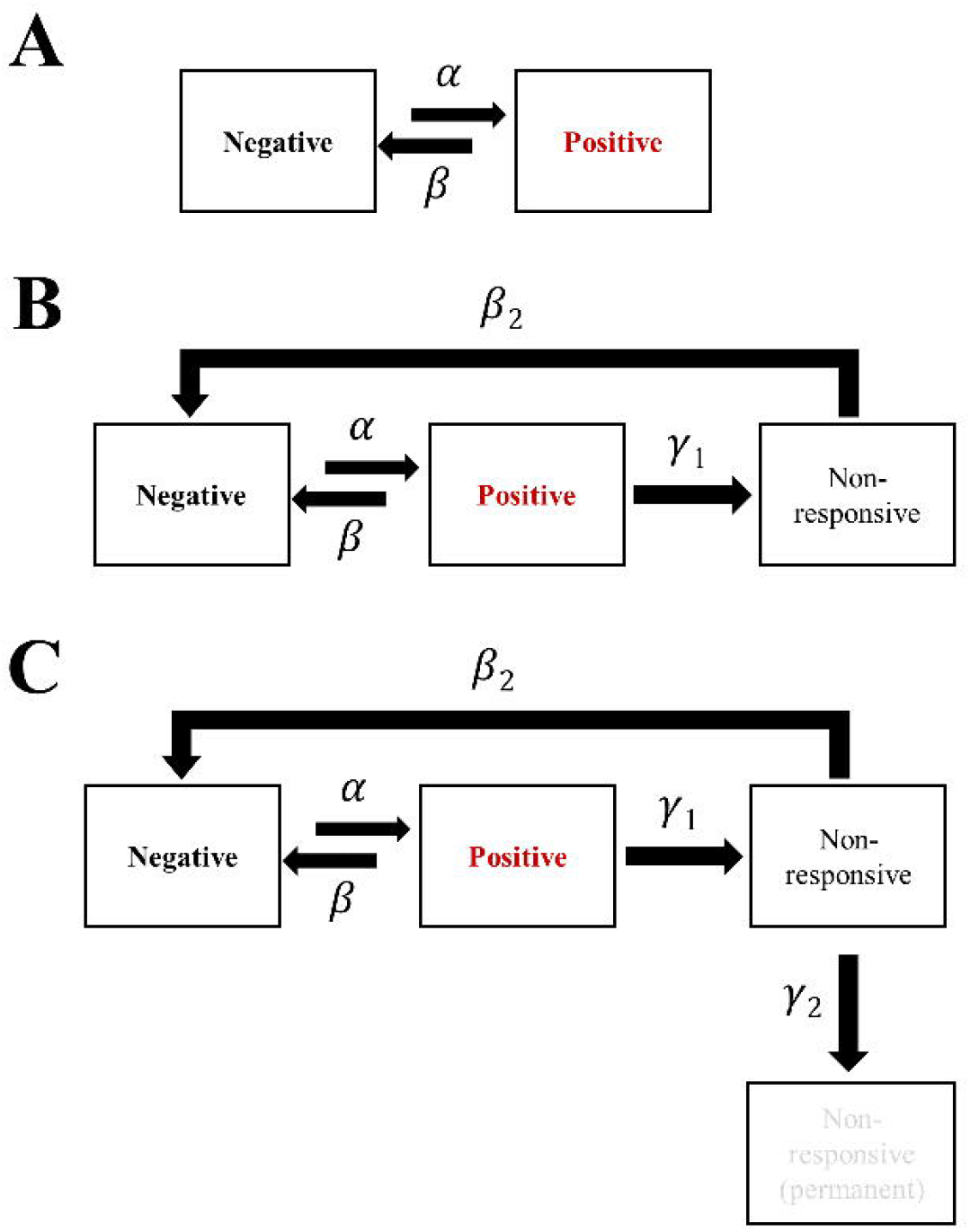
Mathematical modelling with 3 or 4 non-responsive states (NoRM) can describe hyporesponsiveness. A. Schematic representation of states and constants used in a 2-state model. Macrophage can be in a negative or positive state of making an inflammatory response protein. B. Schematic representation of states and constants used in the NoRM mathematical model. Macrophage can be in a negative, positive, non-responsive (NRS). C. Same as B but with inclusion of a 4th non-responsive permanent (NRPS) state.

We termed our overall modelling approach the “non-responsive macrophage” (NoRM) model (**Figure 4**). We stress that the purpose of these models was not to predict detailed physiological transitions or identify mechanisms. Rather, they offered a framework within which to interpret our empirical datasets and alluded to simple explanations for observed phenomena across a range of experimental conditions. In this context, we note that in the NoRM model, all cells were expected to respond to LPS treatment. This assumption also captured cells that might never respond to LPS by transitioning from positive to non-responsive states almost immediately upon stimulus.

### A 3-state NoRM model is sufficient to explain macrophage hypo-responsiveness

Using rejection sampling, we tested whether the 3- or 4-state NoRM models could independently capture our empirical data for each of the measured proteins. Based on the AIC values comparing model fit to estimated parameters, a 3-state NoRM model is sufficient to explain our empirical data (**Supplementary Figure S3B**). We next compared model outputs for proportion of cells in the positive state over time for the 3-state and 4-state NoRM model both of which predict hypo-responsiveness of the population (**Supplementary Figure S4)**. The output from the models was used to predict the composition of positive, negative, NRS and/or, in the case of the 4-state model, NRPS for each of the four proteins. Based on the estimated parameters, our model predicted that the total proportion of non-responsive cell-states (NRS and/or NRPS) increased post primary LPS stimulus (**Figure 5, Figure S5**) and therefore contributed to the diminished response by the population in the second challenge of LPS for all proteins (**Figure S4**) except NOS2 in both cellular models (**Figure S4B** and as seen in the empirical data shown in **Figure 3A**).

**Figure 5:**
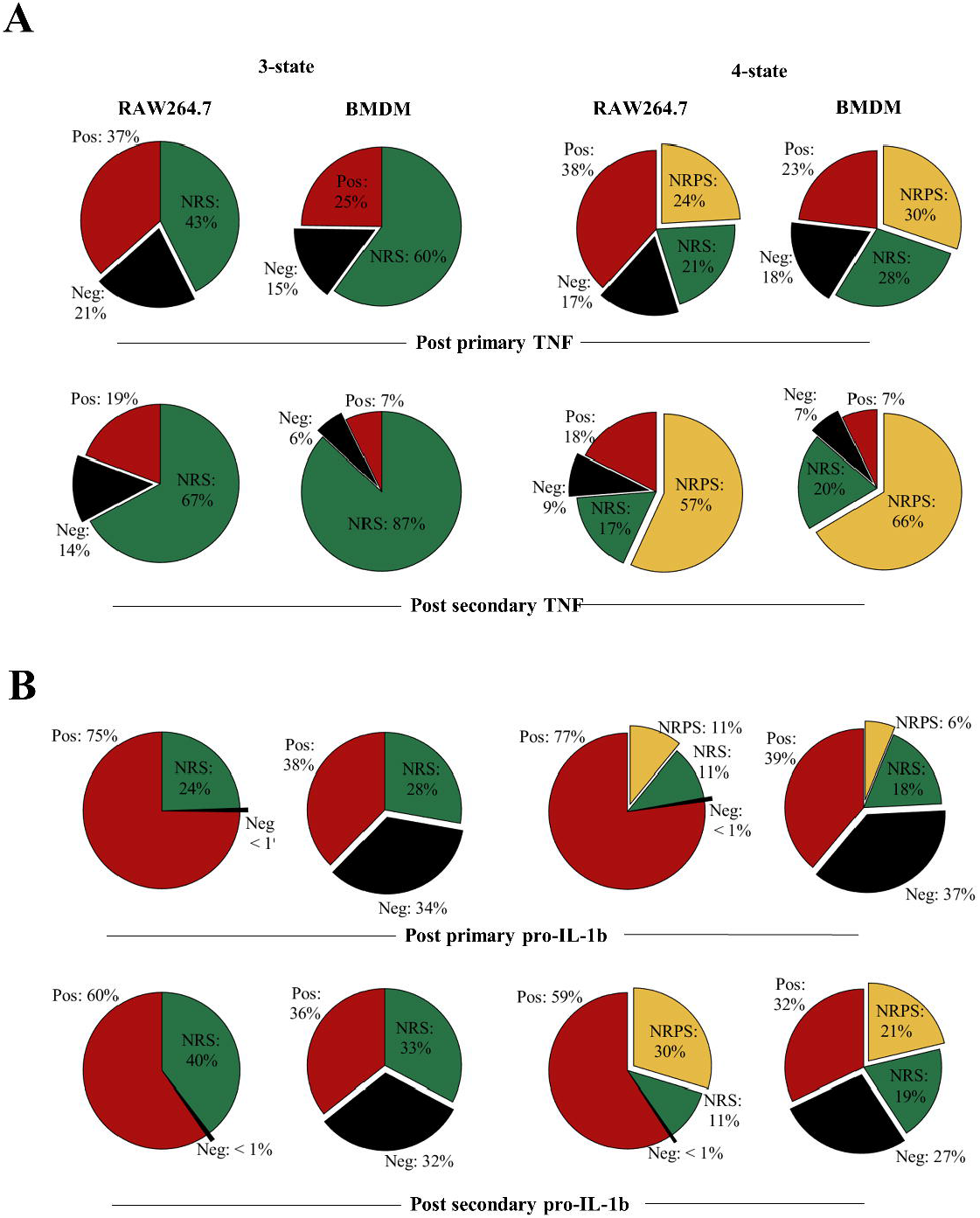
Transitions between distinct non-responding macrophage subsets underpin responses to LPS. A. Overall cell-state compositions for TNF based on the NoRM model prediction when 3-states (ie γ2=0) or 4-states are modelled post in silico stimulation with a single dose of LPS of 1000ng/ml, 12 hours BMDM or 16 hours RAW264.7) and two doses of LPS (1000ng/ml 0-24 hours + 1000ng/ml, 12 hours BMDM or 16 hours RAW264.7). B. Same as above but for pro-IL-1β.

Upon comparing the *in-silico* 3-cell-state composition for each of the inflammatory protein, differences and similarities between BMDM and RAW264.7 cells were visible at 12 and 16 hours of primary *in-silico* stimulus between TNF, IL-6, pro-IL1β and NOS2 (**Figure 5, Supplementary Figure S5, 3-state**). The stimulus length was interpreted based on the empirical results in **Figures 1, 2** and **3**. For TNF (3-state, **Figure 5A**), BMDMs had a higher frequency of cells in the NRS than RAW264.7 cells (60% versus 43%) but despite this both maintained a proportion of cells in the negative state (15% versus 21%). This was compatible with the possibility of a fraction of cells remaining negative but capable of responding at later timepoints. In the case of TNF, when the negative state to NRS ratio is calculated in BMDMs, about 1 in 2 of phenotypically negative cells (for a single protein) can respond to LPS while in RAW264.7 cells this decreases to 1 in 4 suggesting that RAW264.7 cells may show greater sensitivity to becoming TNF+ later into the stimulus. In a similar but with opposite manner, for pro-IL-1β RAW264.7 cells have negative to NRS ratio less than 1:24 at 16 hours (3-state, **Figure 5B**) while the same ratio is greater than 1 in BMDMs. While this could be due to the large difference in positive pro-IL-1β cells in RAW264.7 versus BMDM, it suggested that up to 34% BMDMs remained antigen (LPS) responsive. Interestingly, the 3-state NoRM model suggested similar IL-6 dynamics showing that BMDMs maintained a large negative to NRS ratio after primary (54% negative to 41% NRS) and secondary (52% negative to 43% NRS) LPS stimulation (**Supplementary Figure S5**). The above observations demonstrated differences between the two cellular models and their responsiveness to LPS.

While the 3-state model was sufficient to explain our experimental data points, it did not differentiate between temporary non-responsiveness and permanent epigenetic cessation of activity (Seeley and Ghosh, 2017). To explore how these states might vary between proteins and cell types, we also analysed the 4-state representation of the NoRM model (**Figure 5, supplementary Figure S5, 3-state**). The NRS to NRPS ratio varied greatly between different proteins after the primary (12 hour for BMDM and 16 hour for RAW264.7) and secondary (24hr primary + 12hr/16hr secondary for BMDM and RAW264.7 respectively) dose of LPS (*in silico*). TNF NRPS frequencies were almost 3 times higher than pro-IL-1β in both cellular models. On the other hand, IL-6 NRPS frequency was comparable to TNF NRPS frequency in RAW264.7 but lower (1:3) in BMDMs. Further, in RAW264.7 cells, NOS2 NRPS frequency was less than 1% even after secondary stimulus while in BMDMs this was 5%. The increase in NRPS for any single protein over a subsequent stimulation, however, was consistent for all proteins. This suggested that while some proteins switched off faster in single cells over a course of stimulation, if stimulation remained (i.e. until LPS>0 in the model) the system would progress to all cells becoming non-responsive permanently (NRPS) given γ_2_≠0. Furthermore over the course primary/secondary stimulus (within our modelling timeframe), BMDMs consistently comprised of fewer cells in the NRPS than RAW264.7 cells for TNF, pro-IL-1β and IL-6 with the exception of NOS2 where BMDM communities had higher NRPS frequency.

Taken together, analysis of the NoRM model demonstrated that the existence of one non-responsive macrophage cell state is necessary to explain the observed empirical data. However, a 4-state model including distinct reversible and permanently non-responsive macrophage cell states was also compatible with the empirical data and captured differences between a model macrophage cell line (RAW264.7 cells) and primary macrophages (BMDMs).

## Discussion

Heterogeneity is a hallmark of immune cell populations (Sallusto and Lanzavecchia, 2009;Satija and Shalek, 2014;Guilliams et al., 2018;Papalexi and Satija, 2018). Understanding the mechanisms driving this heterogeneity can reveal how it can be modulated to prevent immunopathology or boost immunity when necessary (Gogos et al., 2000;Rittirsch et al., 2008;Hotchkiss et al., 2013;Davenport et al., 2016). In this context, macrophages pre-exposed to LPS show a dampened immune response when re-stimulated with LPS. This effect is physiologically relevant in the appearance of an immuno-suppressive phase in sepsis and is associated with increased mortality (Biswas and Lopez-Collazo, 2009). Our results reveal that analysis of only a small number of pro-inflammatory proteins combined with simple mathematical models can provide powerful insight into the functional relevance of macrophage AIH. We show how single cells show considerable heterogeneity in production and co-expression of TNF, IL-1β, IL-6, or NOS2, underpinned by functionally distinct non-responsive states. It is of note that although both AIH and non-responsiveness are concepts that have been long used in T cell responses (Schwartz, 2003;Zhu and Paul, 2010), their application and understanding in macrophage responses is profoundly lacking. Our results suggest that heterogeneity in terms of community composition is maintained in hypo-responsive macrophage communities despite the overall lower response and that, at least for a subpopulation of cells, the apparent lack of response is reversible. In our study we measured protein levels of selected key inflammatory mediators using BFA and obtained consistent results in two different macrophage models. Further studies, using single cell proteomics and transcriptomics can be used to define the key molecular features of non-responsive macrophage subsets within a population responding to antigen *in vitro* and *in vivo* and the molecular regulators driving transitions between responding and non-responding macrophage communities. Using ultra-pure LPS in these studies will allow for accurate determination of quantitative effects of AIH. Identifying molecular mechanisms that favor or repress the generation of permanently non-responsive macrophage population can have far-reaching implications for treatment and understanding of infectious, inflammatory, and autoimmune diseases. Similarly, with regards to the mathematical modelling approach, we note that while generating accurate predictions of temporal evolution of protein positivity was not a primary purpose of the NoRM model, it provides a framework to which linear or non-linear constraints to μ (LPS co-efficient) and δ (LPS decay) can be added to model generalised protein positivity at phenomenological levels. This would allow to model primary and secondary effects at objective level generating simple parameters to test in laboratory experiments.

Both our empirical and theoretical analysis of macrophage AIH highlighted differences between RAW264.7 cells and primary BMDMs, for example with regards to kinetics of activation. This is in agreement with proteomics and transcriptomics studies comparing BMDMs with macrophage-like cell lines (Guo et al., 2015;Levenson et al., 2018) indicating differential kinetics and maximum magnitude of responses. Differences in pre-existing genetic heterogeneity and signaling and transcriptional networks between the two cell types are likely sources for these differences. However, there also notable similarities between the two cellular models. For example, in both models macrophages that are challenged with LPS for a second time respond through distinctly different community composition trajectories than those observed in cells that respond to LPS for the first time. Similarly, in the 4-state NoRM model, for both cell types LPS-induced hypo-responsiveness post-secondary challenge is associated with an increase in NRPS. This concurs with reports highlighting that non-reversible mechanisms leading to permanent changes within the cell, such as chromatin remodeling, are critical for induction of endotoxin tolerance (Seeley and Ghosh, 2017). In a biological context, the NRS can be considered as arising from sufficient but temporary effects such as post-transcriptional attenuation of the TLR4 pathway and/or miRNA induced, while the NRPS might represent longer heritable epigenetic modifications (Nomura et al., 2000;Chan et al., 2005;Quinn et al., 2012;Seeley and Ghosh, 2017;Vergadi et al., 2018). Overall, it is important to note the value of our approach in revealing cellular ecology and community dynamics aspects that align with molecular and phenotypic insights into training of macrophages (Saeed et al., 2014;Netea et al., 2016).

Variability in gene expression in eukaryotic cells (McAdams and Arkin, 1997;Elowitz et al., 2002;Paulsson, 2004) often has phenotypic consequences (Blake et al., 2006;Eldar and Elowitz, 2010). Innate immune response to stimulus has been shown to be heterogeneous in mammalian immune cells (Shalek et al., 2013;Satija and Shalek, 2014). This is most notable in the high transcriptional variability of cytokines, such as TNF, IL-1β, and IL-6, and their receptors upon stimulus in LPS stimulated phagocytes (Hagai et al., 2018). *In vivo*, the source of macrophage population heterogeneity could be driven by developmental, tissue or niche, and activation associated factors. Furthermore, it can be amplified or suppressed through interaction with other immune or non-immune cells (Yao et al., 2018). Our study explored macrophage AIH exclusively *in vitro* using relatively homogeneous starting cell populations to concentrate on cell-intrinsic mechanisms. Nevertheless, it is likely that our theoretical model only partially captures population heterogeneity occurring in more complex macrophage populations or *in vivo*. However, we speculate that the key concepts revealed by our findings including AIH dose dependence, existence of reversible and permanently non-responsive states, and a critical role for transitions between these states as determinants of macrophage function will be relevant to a broad range of pathophysiological contexts in the immune system.

## 2 Conflict of Interest

The authors declare that the research was conducted in the absence of any commercial or financial relationships that could be construed as a potential conflict of interest.

## 3 Author Contributions

D.L. and J.W.P. conceived and supervised the project. S.D., D.B., J.W.P., and D.L. designed experiments and analysis pipelines. S.D. performed and analysed experiments, developed the mathematical model. All authors co-wrote the manuscript.

## 4 Funding

The study was funded by the “Combating Infectious Disease: Computational Approaches in Translational Science” Wellcome Trust PhD programme (WT095024MA).

## 5 Acknowledgments

We thank staff at the Imaging and Cytometry Lab in the University of York Bioscience Technology Facility for technical support and advice.

## Notes

### Competing Interest Statement

The authors have declared no competing interest.

