## Supplementary for "Mathematical modelling of activation-induced heterogeneity reveals cell state transitions underpinning macrophage responses to LPS"

### Supplementary Material

#### 1 MODELLING PROCESS

The supplementary material describes the modelling process in detail. In particular, it describes simpler models that were tested to arrive at our proposed model for macrophage activation.

##### 1.1 Positive-state model definition

The response to LPS in the *in silico* cell environment was modelled as a direct positive effect on the forward rate that describes the transition of a cell to the P state such that

$$\alpha_{LPS} = \alpha(1 + \mu L) \tag{S1}$$

where $\alpha_{LPS}$ is the overall forward rate which takes into account the LPS in the environment, $\alpha$ is the forward rate in the absence of LPS, $L$ is the concentration of LPS in the environment and $\mu$ is a constant that describes the magnitude of the response to LPS concentration. The linear assumption in **equation S1** is used for simplicity to induce plausible local LPS dynamics at the cost of introducing a single unknown, $\mu$.

Upon describing the above model as an ODE, we have

$$\frac{dP}{dt} = N\alpha(1 + \mu L) - \beta P \tag{S2}$$

where $\frac{dP}{dt}$ represents rate of change in the number of cells in the positive state with respect to time, $P, N$ are the number of *in silico* cells in the positive and negative state respectively, $\beta$ is the rate at which cells in the positive state change to negative state.

We model the decay of LPS concentration, $L$ as a simple first order exponential decay in continuous time. This can then be expressed as

$$L(t) = L_0 e^{-\delta t} \tag{S3}$$

where $L_0$ and $L(t)$ represent the concentration of LPS at time zero and $t$ respectively, and $\delta$ is the constant LPS decay rate.
At a time dependent quasi-equilibrium, where the dynamics of $L$ are assumed to occur on a slower time scale than those of $P$ and $N$, **equation S2** can be used to write

$$P^*_{quasi} = \frac{N\alpha(1 + L_0 e^{-\delta t})}{\beta} \tag{S4}$$

where rate of change of $P$, $\frac{dP}{dt} = 0$ and $P^*_{quasi}$ represents the number of *in silico* cells in the positive-state at quasi-equilibrium. It can be inferred from **equation S4** that in the absence of LPS in the environment,

$$P^* = \frac{N\alpha}{\beta} \tag{S5}$$

which is the equilibrium ($P^*$) dynamics for the simple case where 2 species switch between each other with rates $\alpha$ and $\beta$ respectively.

Further, **equation S2** can be re-written as

$$\frac{dP}{dt} = (T - P)\alpha(1 + \mu L) - \beta P \tag{S6}$$

where T is total number of cells since $N + P = T$ at any given time.
Equation S6 can be solved exactly to the following closed form equation given the initial condition $P(0) = 0$

$$P(t) = e^{\frac{L_0 \alpha \mu e^{-\delta t}}{\delta} - t(\alpha+\beta)} \int_0^t T\alpha e^{\alpha t + \beta t - (L_0 \alpha \mu e^{-\delta t})/\delta} (L_0 \mu e^{-\delta t + 1}) dt \tag{S7}$$

While **equation S7** can model the fraction of macrophages responding to a primary LPS stimulation, it fails to explain the effect of second dose of LPS wherein a smaller fraction of the population responds unless the value of $\alpha$ is modified for the population. This implies that the LPS response of the whole population changes upon secondary stimulus and is incompatible with experimental results, at the transcript level, cytokines are expressed bi-modally (Shalek, 2013 #49).

#### 1.2 Non-responsive state model ("3-state model")

We then reasoned if the inclusion of a single state (phenotype) could explain macrophage hyporesponsiveness with respect to any one protein.

To this effect, we first excluded the LPS-induced effect on $\alpha$ and explored simpler linear explanations of what we observe by considering the following equations

$$\frac{dN}{dt} = -N\alpha_{LPS} + \beta P \tag{S8}$$

$$\frac{dP}{dt} = N\alpha_{LPS} - \beta P - \gamma_1 P \tag{S9}$$

$$\frac{dNR}{dt} = \gamma_1 P - \beta_2 NR \tag{S10}$$

where $NR$ represents the non-responsive state and $\gamma_1$ represents the rate at which positive cells change to a non-responsive state ($NR$, S10) while $\beta_2$ rate at which non-responsive cells become negative cells ($N$, S8) and $\alpha_{LPS}$ is LPS dependent (S1).

#### 1.3 Non-responsive state model ("4-state model")

We then asked whether the inclusion of an additional state (a non-responsive permanent state or NRPS) can resolve empirical hyporesponsiveness better. NRPS state is modelled as an end-point state in the model with the rate of change equation for NRPS state and modifying equation S10 as following

$$\frac{dNRPS}{dt} = \gamma_2 NR \tag{S11}$$

$$\frac{dNR}{dt} = \gamma_1 P - \beta_2 NR - \gamma_2 NR \tag{S12}$$

where $\gamma_2$ is the rate at which cells become permanently non-responsive.

Due to variability in experimental data (see **Figure 2** versus **Figure 3** for M/1000 at 4hr+4hr BFA) we further modelled the above ODE model using the Doob-Gillespie algorithm to account for interpreting variability as stochastic noise (NoRM model). We first checked if the NoRM model and ODE model agree in terms of the latter representing a mean-field for the stochastic model (**Figure S1A**). Using rejection sampling (details in Methods section) we checked which of the above models were consistent with our experimental data (**Figure S1B**) .

### 2 SUPPLEMENTARY FIGURES

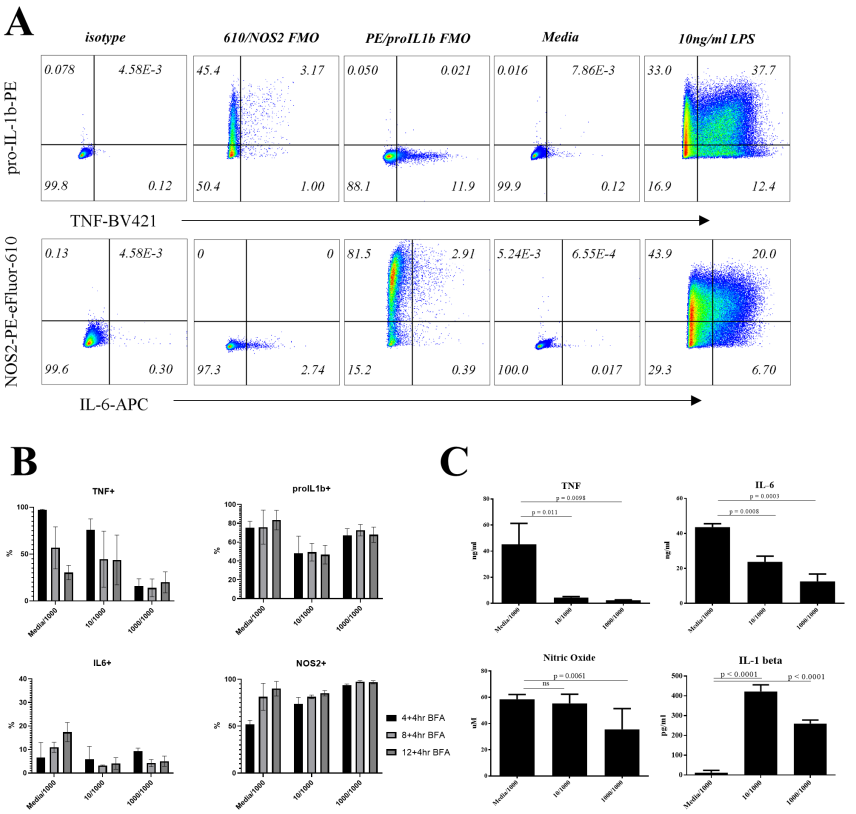


**Figure S1. Altered cytokine production kinetics in RAW264.7 macrophages responding to a second LPS challenge** A) Bi-plots describing strategy for gating for positive and negative fractions for TNF, pro-IL1$\beta$, IL-6 and NOS2 positive cells throughout the experiments shown here for BMDMs as a representation. Gating was adjusted based on pooled isotype, and re-confirmed using fluorescent minus one controls (FMOs for PE and PE-eFluor 610) with controls for media only treatment. Finally a representative graph showing BMDMs treated with 10ng/ml LPS to show staining example and positive fractions. B) Percentages for TNF, pro-IL1$\beta$, IL-6 and NOS2 positive cells at the indicated timepoints post LPS challenge (1000ng/ml) in RAW264.7 cells pre-treated for 24 hours with either media (Media/1000), or 10ng/ml LPS (10/1000), or 1000ng/ml LPS (1000/1000). C) TNF, IL-6, IL-1$\beta$ and Nitric Oxide (NO) cumulative secreted levels at 24 hours post-secondary stimulation with 1000ng/ml LPS in macrophages pre-treated for 24 hours with either media (Media/1000), or 10ng/ml LPS (10/1000), or 1000ng/ml LPS (1000/1000). n=3-6 per treatment group.

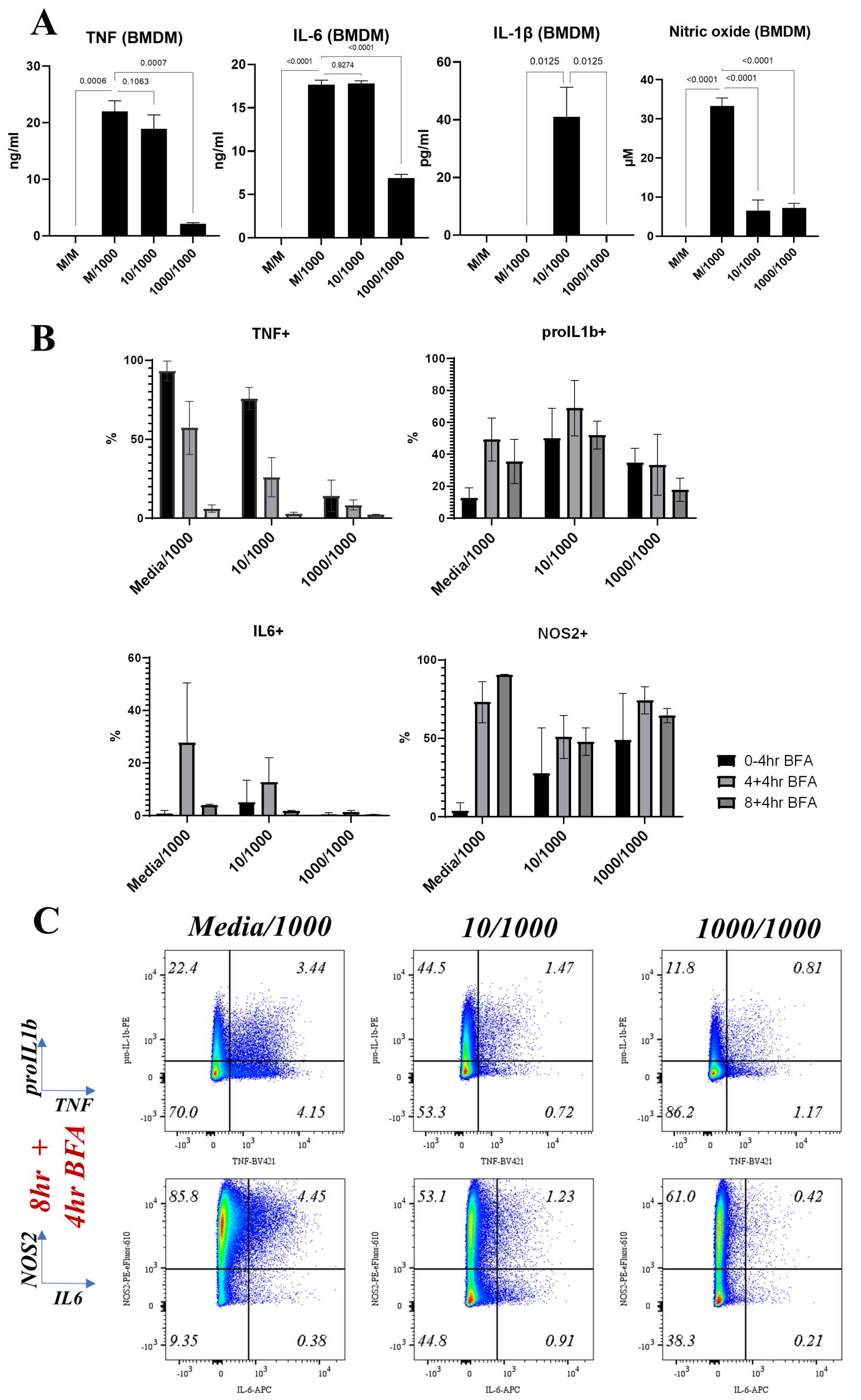


**Figure S2. BMDMs show a clear hyporesponsive phenotype by flow** A) TNF, IL-6, IL-1$\beta$ and Nitric Oxide (NO) cumulative secreted levels at 24 hours post-secondary stimulation with 1000ng/ml LPS in macrophages pre-treated for 24 hours with either media (M/1000), or 10ng/ml LPS (10/1000), or 1000ng/ml LPS (1000/1000). n=4 per treatment group. B) Percentages for TNF, pro-IL1$\beta$, IL-6 and NOS2 positive cells at the indicated timepoints post LPS challenge (1000ng/ml) in BMDMs pre-treated for 24 hours with either media (Media/1000), or 10ng/ml LPS (10/1000), or 1000ng/ml LPS (1000/1000). C) Bi-plots showing TNF, pro-IL-1$\beta$ and IL-6, NOS2 expression at indicated timepoints post LPS challenge (1000ng/ml) in BMDMs pre-treated for 24 hours with either media (Media/1000), or 10ng/ml LPS (10/1000), or 1000ng/ml LPS (1000/1000). Cells were pre-gated on Live/Singlets/Forward and Side Scatter/CD11b+F4/80+. Plots are representative of 3 independent experiments.

**A**

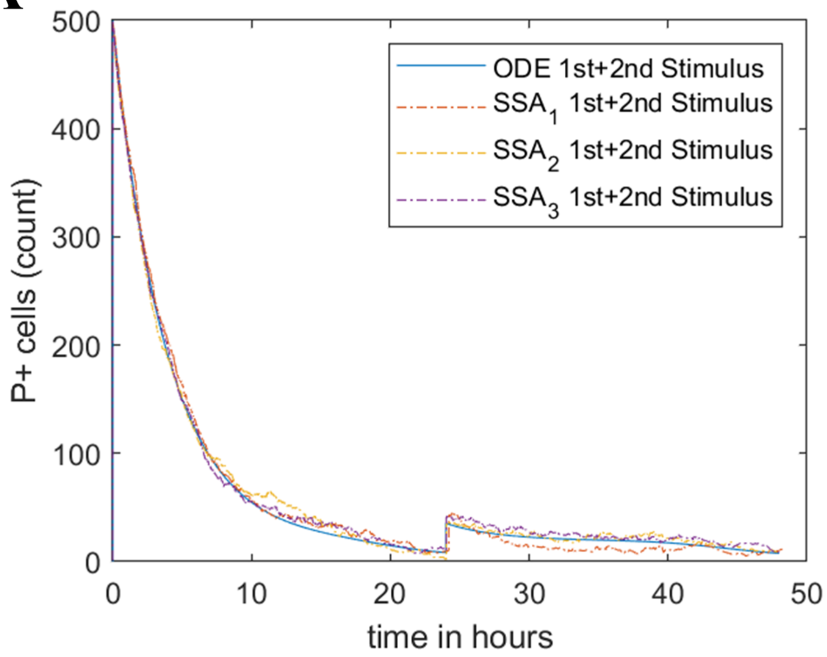


**B**

| | 2-state + LPS | 3-state + LPS | 4-state + LPS |
|---|---|---|---|
| | | BMDMs | |
| TNF | X | -24.4 | -23.1 |
| IL-6 | X | -33.5 | -31.5 |
| NOS2 | X | -16 | -13.4 |
| pro-IL-1b | X | -20.9 | -18.9 |
| | | RAW264.7 | |
| TNF | X | -23.9 | -22.7 |
| IL-6 | X | -46.9 | -45.9 |
| NOS2 | X | -26.2 | -23.8 |
| pro-IL-1b | X | -31.5 | -30.2 |

**Figure S3. Mathematical modelling with 3 or 4 non-responsive states can describe hyporesponsiveness** A) Doob-Gillespie algorithm for NoRM model simulations (n=3) are overlaid on the mean-field ODE based non-responsive model with 3-states. Time=0hr represents first in silico LPS stimulation and time=24hr represents second LPS stimulation. Arbitrary parameters were chosen for proof of concept B) Table depicting average Akaike Information Criterion (AIC) values obtained for top 50 estimated parameter sets for 3-state and 4-state NoRM models independently per protein. AIC was not estimated for 2-state model.

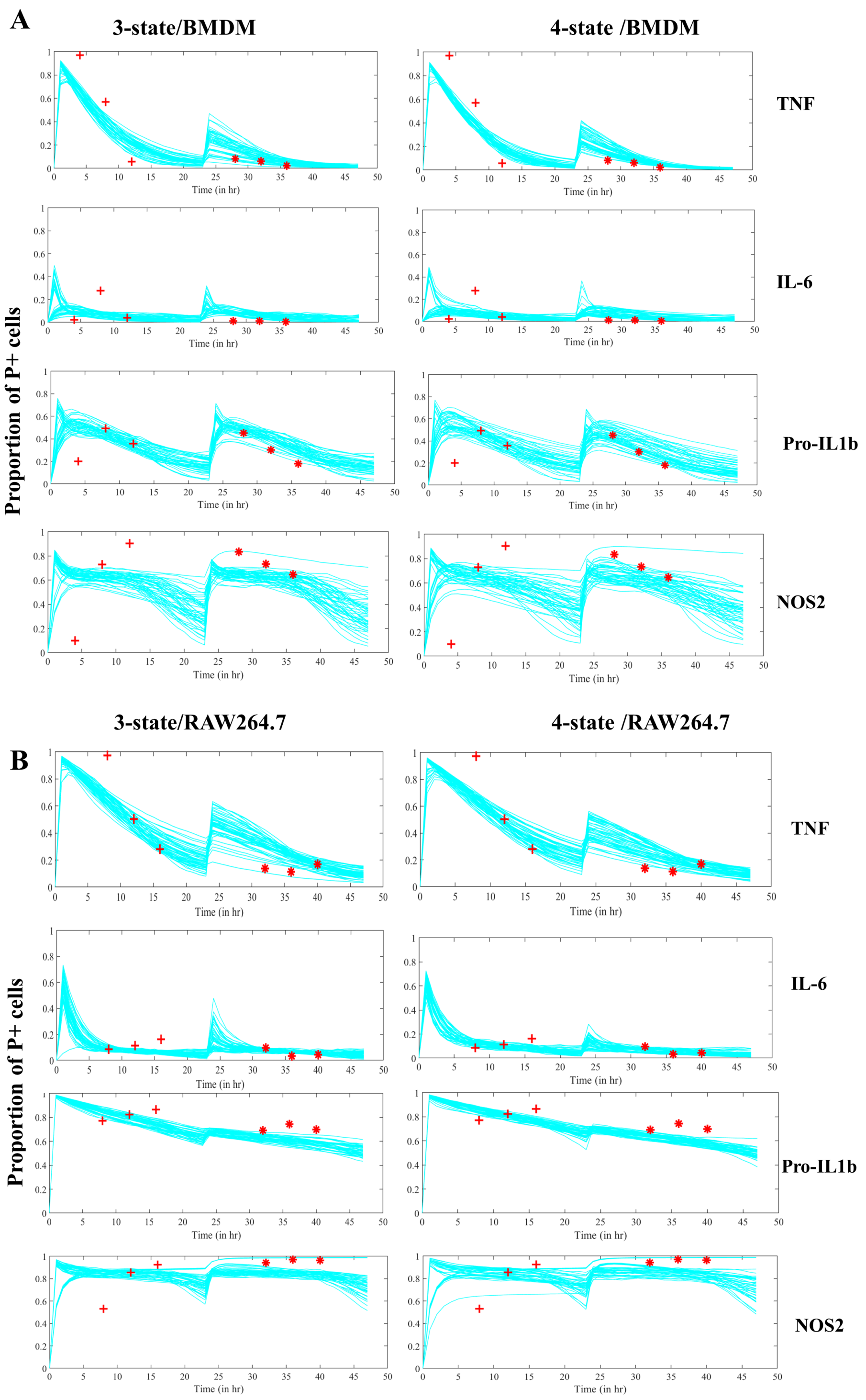


**Figure S4. Estimated trajectories of NoRM 3 and 4-state model** A) Two LPS stimulations are simulated using the NoRM models (3-state left, 4-state right) using top 50 estimated parameter sets based on experimental timepoints marked in red. Time=0hr represents first *in silico* LPS stimulation and time=24hr represents second LPS stimulation for BMDMs B) Same as above but for RAW264.7 cells.

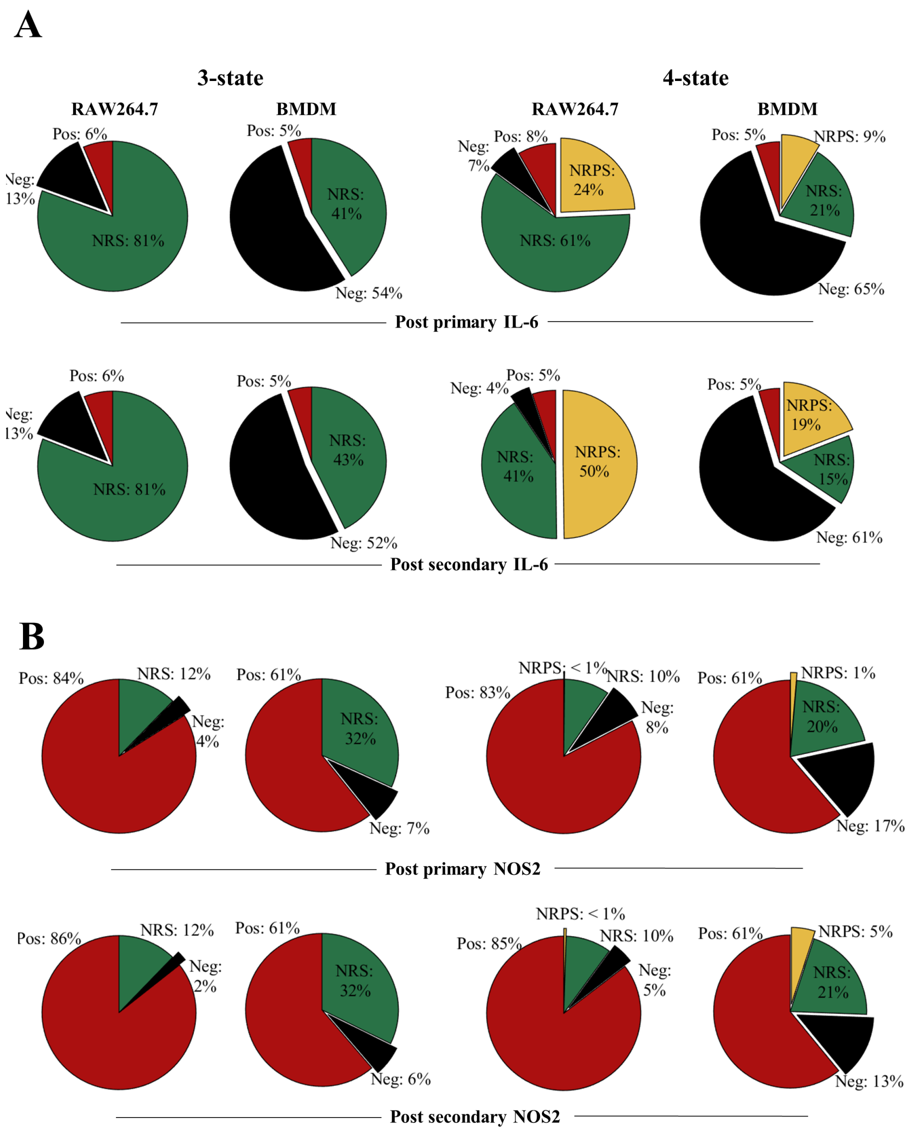


**Figure S5. Transitions between distinct non-responding macrophage subsets underpin responses to LPS** A) Overall cell-state compositions for pro-IL1b based on the NoRM model prediction when 3-states (ie $\gamma_2 = 0$) or 4-states are modelled post *in silico* stimulation with a single dose of LPS of 1000ng/ml, 12 hours BMDM or 16 hours RAW264.7) and two doses of LPS (1000ng/ml 0-24 hours and 1000ng/ml , 12 hours BMDM or 16 hours RAW264.7). B) Same as above but for NOS2.
